# Spatial control over near-critical-point operation ensures fidelity of ParAB*S*-mediated bacterial genome segregation

**DOI:** 10.1101/2020.04.26.062497

**Authors:** Longhua Hu, Jérôme Rech, Jean-Yves Bouet, Jian Liu

## Abstract

In bacteria, most low-copy-number plasmid and chromosomally encoded partition systems belong to the tripartite ParAB*S* partition machinery. Despite the importance in genetic inheritance, the mechanisms of ParAB*S*-mediated genome partition are not well understood. Combining theory and experiment, we provided evidences that the ParAB*S* system – partitioning via the ParA gradient-based Brownian ratcheting – operates near a critical point *in vivo*. This near-critical-point operation adapts the segregation distance of replicated plasmids to the half-length of the elongating nucleoid, ensuring both cell halves to inherit one copy of the plasmids. Further, we demonstrated that the plasmid localizes the cytoplasmic ParA to buffer the partition fidelity against the large cell-to-cell fluctuations in ParA level. Thus, the spatial control over the near-critical-point operation not only ensures both sensitive adaption and robust execution of partitioning, but sheds light on the fundamental question in cell biology: How do cells faithfully measure cellular-scale distance by only using molecular-scale interactions?

## INTRODUCTION

Cellular processes must establish the right operating point in a very large parameter space – that allows robust execution of biological function and simultaneously sensitive adaptation to environmental cues. What is the character of this right operating point? How do cells find and maintain it? It is postulated that all living systems operate near the edge of phase transition (Mora and Bialek 2011), *i.e.*, near a critical point where the system is halfway between two phases in its parameter space. Operating near such a tipping point allows the system to sensitively adapt to changes. While increasing evidences support this notion (Hesse and Gross 2014; Krotov et al. 2014; Mojtahedi et al. 2016; Tetzlaff et al. 2010), they are largely statistical inference from experimental data and lack the physical mechanisms that give rise to the near-tipping-point behavior. Crucially, it is unclear how cellular processes maintain the robustness of operating near the critical point, in the presence of ever-lasting noises (*e.g.*, the fluctuations in gene expression at a single-cell level (Elowitz et al. 2002; Kærn et al. 2005; Newman et al. 2006)). From this perspective, we set out to examine the physical mechanism of bacterial DNA segregation with the emphasis on how the operating point of DNA partition machinery is controlled to maximize the partition fidelity. We exploited low-copy-number plasmid partition in bacteria as the model system by combining theoretical modeling with experimental testing.

Segregating replicated genomes before cell division is essential to ensure faithful genetic inheritance. Despite its simple form, partitioning of low-copy-number plasmids in bacteria is robust with an extremely low error rate (less than 0.1% per generation) (Nordström and Austin 1989), and provides a tractable paradigm to understand fundamental principles of genome segregation (Bouet and Funnell 2019). Most low-copy-number plasmids are actively partitioned – by a conserved tripartite ParAB*S* system – along the nucleoid, a rod-like structure consisting primarily of condensed chromosomal DNA. While cargo trafficking *in vivo* typically utilizes cytoskeletal filament or motor protein-based mechanisms, the ParAB*S* machinery utilizes none of the conventional mechanisms for partitioning (Hatano and Niki 2010; Lim et al. 2014; Vecchiarelli, Hwang, and Mizuuchi 2013; Vecchiarelli, Neuman, and Mizuuchi 2014). How the ParAB*S* system drives genome partitioning has puzzled the field since its first postulation in the replicon theory (Jacob, Brenner, and Cuzin 1963). The key elements of ParAB*S* system are as follows (Bouet and Funnell 2019): ParA is an ATPase that binds non-specifically to DNA in nucleoid in an ATP-bound dimeric state. ParB is the adaptor protein. It binds specifically at a centromere-like site *parS* on the plasmid and sequence-nonspecifically around *parS* to form large clusters called partition complexes (PCs). ParB regulates ParA DNA binding by (i) direct interaction that provokes its release from the nucleoid (Vecchiarelli, Hwang, and Mizuuchi 2013), and by (ii) stimulating the ATPase activity that convert ParA in the ADP-bound form that do not bind nucleoid DNA (Bouet et al. 2007; Leonard, Butler, and Löwe 2005). It is not understood how the chemical energy provided by ATP hydrolysis is harnessed to ensure the PC partition fidelity, beside its important implication in separating newly duplicated *parS* sites (Ah-Seng et al. 2013).

Spatial-temporal features of the ParAB*S* system expose some clues of its inner-working. PCs move around, and frequently switch directions over the cell length (Gordon et al. 2004; Hatano and Niki 2010; Niki and Hiraga 1997; Ringgaard et al. 2009; Sengupta et al. 2010; Walter et al. 2017). As the timing of PC replication and segregation is not directly coupled to cell cycle and the nucleoid itself keeps elongating before cell division (Ellasson, Bernander, and Nordström 1996; Helmstetter et al. 1997; Onogi, Miki, and Hiraga 2002), the replicated PCs can be anywhere along the nucleoid length when they start to split apart (Sengupta et al. 2010). Intriguingly, the replicated PCs always first move apart persistently and then position themselves with the separation being ∼ half of the cell length (Gordon et al. 2004; Hatano and Niki 2010; Niki and Hiraga 1997, 1999; Sengupta et al. 2010). While segregating by half of the cell length ensures the partition fidelity by always positioning the replicated PCs in the different cell halves, it precipitates the following questions. First, given that the PCs locally interact with the nucleoid, which elongates in proportion to the cell length (Bakshi et al. 2014), the question becomes: How do the PCs have the global “view” of, and adapt their separation to, the length of the elongating nucleoid and, ultimately, the half of the cell length? Second, what ensures the partition robustness in the presence of ever-lasting noises, *e.g.*, the fluctuations in protein levels? Addressing these questions lay at the heart of one of the fundamental questions in cell biology: How do cells faithfully measure cellular-scale distance by only using molecular-scale interactions?

We previously established the ParA protein gradient-based Brownian ratchet model – a new mechanism of processive cargo transport, without resorting to filament nor motor proteins (Hu et al. 2015, 2017b). With multiple ParA-ParB bonds tethering a *parS-*coated cargo to a DNA carpet, ParB-stimulated bond dissociation triggers the release of the ParA from the DNA carpet, the randomness of which results in a force imbalance that drags the cargo forward. Critically, the time delay in resetting ParA DNA-binding affinity generates a ParA-depletion zone behind the forward moving cargo (Hu et al. 2015; Hwang et al. 2013; Vecchiarelli, Neuman, and Mizuuchi 2014), perpetuating the asymmetry and persistent movement. As such, the ParAB*S* system can work as a Brownian ratchet: the ParB-bound cargo “self-drives” by both creating and following a ParA gradient over the DNA. This protein gradient-based Brownian ratchet model provides a conceptual framework that allowed us to explain – for the first time in a coherent manner – the diverse motility patterns of PCs evidenced *in vivo* (Hu et al. 2017a) and starts to gain support from *in vivo* experiments (Debaugny et al. 2018; Le Gall et al. 2016; Schumacher et al. 2017; Uhía et al. 2018). However, our effort so far focused on a highly simplified picture, it is unclear 1) whether and how this Brownian ratchet mechanism can adapt the plasmid segregation distance to the length of an elongating nucleoid, and 2) how the PC partition ensures its fidelity against cellular noises.

Here, we show that 1) this ParA gradient-based Brownian ratcheting of bacterial low-copy-number plasmid partitioning operates near a critical point *in vivo*, and 2) the spatial controls over the near-critical-point operation allow both sensitive adaptation of partition to the nucleoid length and robust execution of partition to buffer against noises.

## RESULTS

### Qualitative model description

We first built the *in vivo* model – to specifically capture how ParA-mediated PC partition responds to the dynamical changes – associated with nucleoid elongation during cell growth. The qualitative model features are described below (Fig. 1), and the quantitative model formulation are provided in MATERIALS and METHODS.

**Figure 1.**
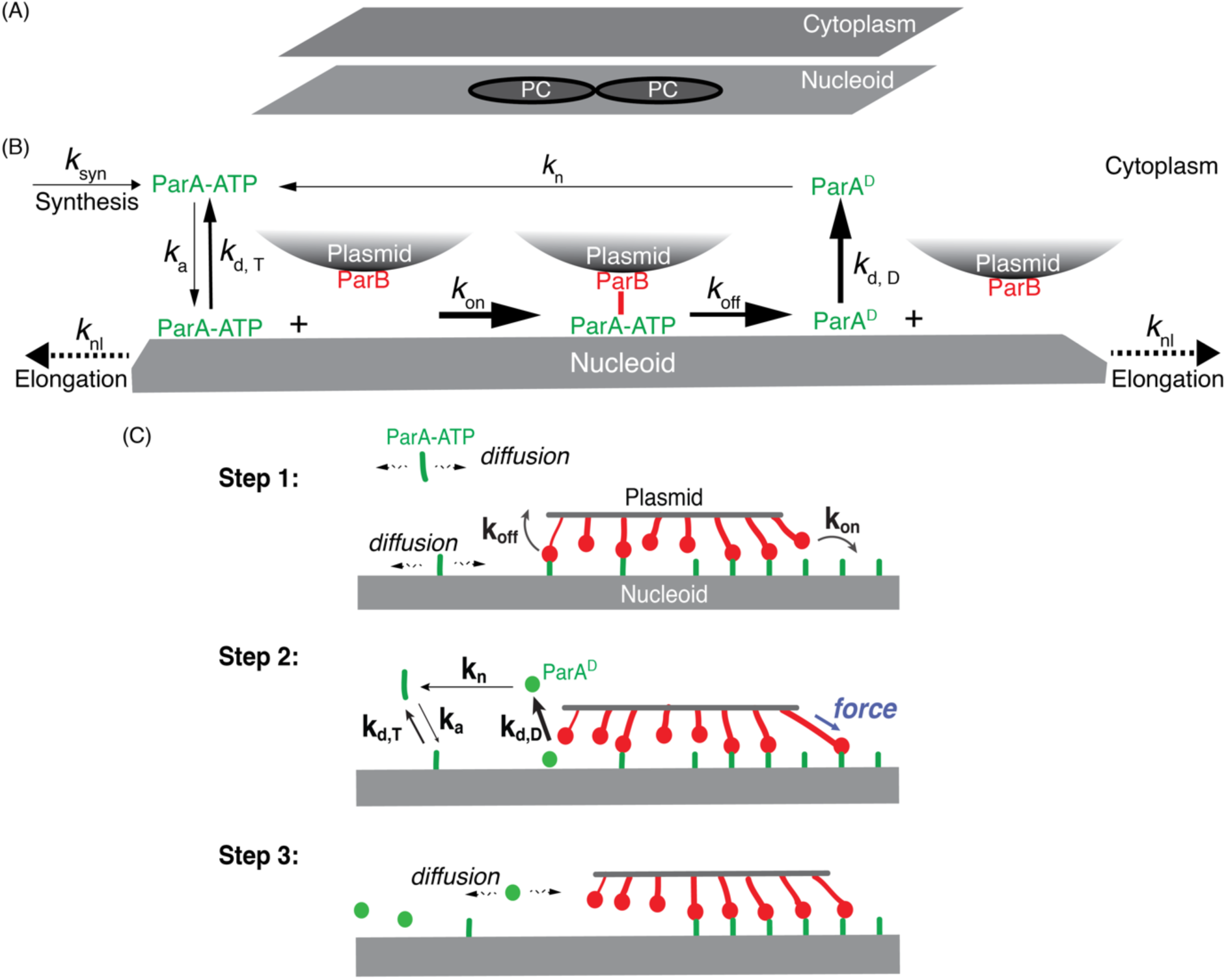
Model description. (A) Model setup. (B) Biochemical scheme of ParABS system. (C) Mechanochemical coupling of ParA-ParB bond formation and dissociation underlies ParA gradient-based Brownian ratcheting. The model parameters are listed in Section I in the supplemental information.

Our model begins with two PCs arranged side-by-side to mimic the replicated PCs and examines the subsequent partition dynamics. As a starting point of the modeling, we depict each PC as a circular disc of ∼ 100 nm in radius (Le Gall et al. 2016; Sanchez et al. 2015), the nucleoid as a flat rectangle, and the cytoplasm as a 2D domain of the same dimension as the nucleoid (Fig. 1A). This is an approximation based on the following considerations. In bacteria such as *E. coli*, the cytoplasm mainly occupies the space of 100 – 200 nm wide between the nucleoid and cell membrane. Because the free ParAs diffuse rapidly in cytoplasm (∼ 3 μm^2^/s)(Surovtsev, Lim, and Jacobs-Wagner 2016), it only takes ParA ∼ 1–10 milliseconds to diffuse across this short distance. Therefore, on the timescale considered (∼ seconds to minutes), the concentration profile of ParA is uniform in the direction vertical to the nucleoid surface, which allows us to simplify the cytoplasm as a flat 2D domain, which serves as a reservoir of free ParAs. Thus, the 2D-domain of cytoplasm in the model represents the effective interface of exchanging free ParAs between the cytoplasm and the nucleoid, whose dimensions was set to be the same.

To capture the dynamic changes associated with nucleoid elongation and cell growth, the model describes two effects. First, with their widths fixed at 1.0 μm, the nucleoid and cytosolic domains elongate at a constant rate (∼ 6-18 nm/min) measured by our experiments (Fig. 1B), in which we imaged HU-mCherry-tagged nucleoid over time in *E. coli* growing in minimal growth medium. The initial nucleoid length is set to be 2 μm, if not otherwise mentioned. Second, the model depicts that concurrent with nucleoid elongation, new ParA molecules are generated to keep ParA concentration constant. This captures the essence of observed auto-regulation of ParA expression (Mori et al. 1989; Yates, Lane, and Biek 1999) and Western-blot measurements showing the constant ParA concentration on population-level (Sanchez et al. 2015).

Accordingly, the model depicts the moving boundary lengthwise and imposes hard-wall boundary condition for ParA and ParB at all edges of the simulation domain. While ParB only localizes to the PCs (Sanchez et al. 2015), ParA can exchange between the nucleoid and the cytoplasm in accordance to the reaction-diffusion scheme (Fig. 1B and Fig. 1C). Specifically, ParA·ATP binds to vacant, unoccupied locations of the nucleoid at a basal rate (Vecchiarelli et al. 2010; Vecchiarelli, Hwang, and Mizuuchi 2013), and can transiently unbind and rebind to adjacent vacant sites via lateral diffusion (Surovtsev, Lim, and Jacobs-Wagner 2016). Upon binding to plasmid-bound ParB, the ParA·ATP no longer diffuses but forms a ParA·ATP-ParB bond, tethering the PC to the nucleoid through the nucleoid-ParA·ATP-ParB-plasmid linkage. For simplicity we refer the entire linkage as the ParA-ParB bond (Fig. 1C).

The ParA-ParB bond formation and the subsequent deformation, similar to deforming a spring, generates a restoring force on the PC (Fig. 1C). The vector sum of many individual ParA-ParB bonds across the PC collectively generates a net force that displaces the PC. The movement of PC in turn changes the bond configurations. When random events (*e.g.*, PC diffusion and stochastic ParA-ParB bond dynamics) break symmetry, the PC moves forward with the ParA-ParB bonds broken at its back (Step 1 in Fig. 1C).

We define the resulting disengaged ParA to be in a distinctive state, ParA^D^ (Fig. 1B). While the model does not specify whether ParA^D^ corresponds to an ATP-bound or ADP-bound state, it recapitulates two key aspects of disengaged ParA. First, ParA^D^ dissociates from the nucleoid faster than the basal turnover rate of ParA·ATP, which reflects the known effect of ParB-mediated stimulation on ParA release from the nucleoid (Vecchiarelli et al. 2010; Vecchiarelli, Hwang, and Mizuuchi 2013). Second, once ParA^D^ dissociates into the cytoplasm, it slowly reverts to the ATP-bound state competent for DNA-binding (Vecchiarelli et al. 2010; Vecchiarelli, Hwang, and Mizuuchi 2013). This time delay results in a ParA-depletion zone trailing behind the moving PC, which subsequently can be refilled by cytosolic and nucleoid-associated ParA·ATP.

As the PC moves forward, the ParBs on the leading edge of the PC continue to establish new bonds with ParA·ATP on unexplored regions of the nucleoid, where the ParA·ATP concentration is higher (Step 2 in Fig. 1C). PC movement therefore maintains the asymmetric ParA concentration gradient that in turn supports further forward movement (Step 3 in Fig. 1C), resulting in a directed and persistent movement. Conceptually, our model is a two-dimensional burnt-bridge Brownian ratchet model (Morozov and Kolomeisky 2007; Saffarian et al. 2006), in which mechanical actions of the multiple bonds not only facilitate the forward cargo movement, but collectively provide tethering that quenches the cargo lateral diffusion. This drives the directed and persistent movement

### ParABS-mediated PC partition operates near a critical point

We formulated the PC partition problem in the framework of dynamical phase transition, which denotes different PC motility modes as distinct states. We defined the order parameter (ψ) – that depicts and distinguishes different states of PC motility – as the PC’s excursion range normalized by half of the nucleoid length. For instance, ψ is ∼ 0 for being stationary and approaches ∼ 2.0 for pole-to-pole oscillation (Fig. 2 – Figure supplement 1). Further, we used the PC segregation distance as a proxy to infer the partition fidelity: In line with our previous work (Hu et al. 2017a), the more the PC segregation deviates from the half length of nucleoid, the more likely will the two PCs end up in the same half of the dividing cell, compromising the partition fidelity.

**Figure 2.**
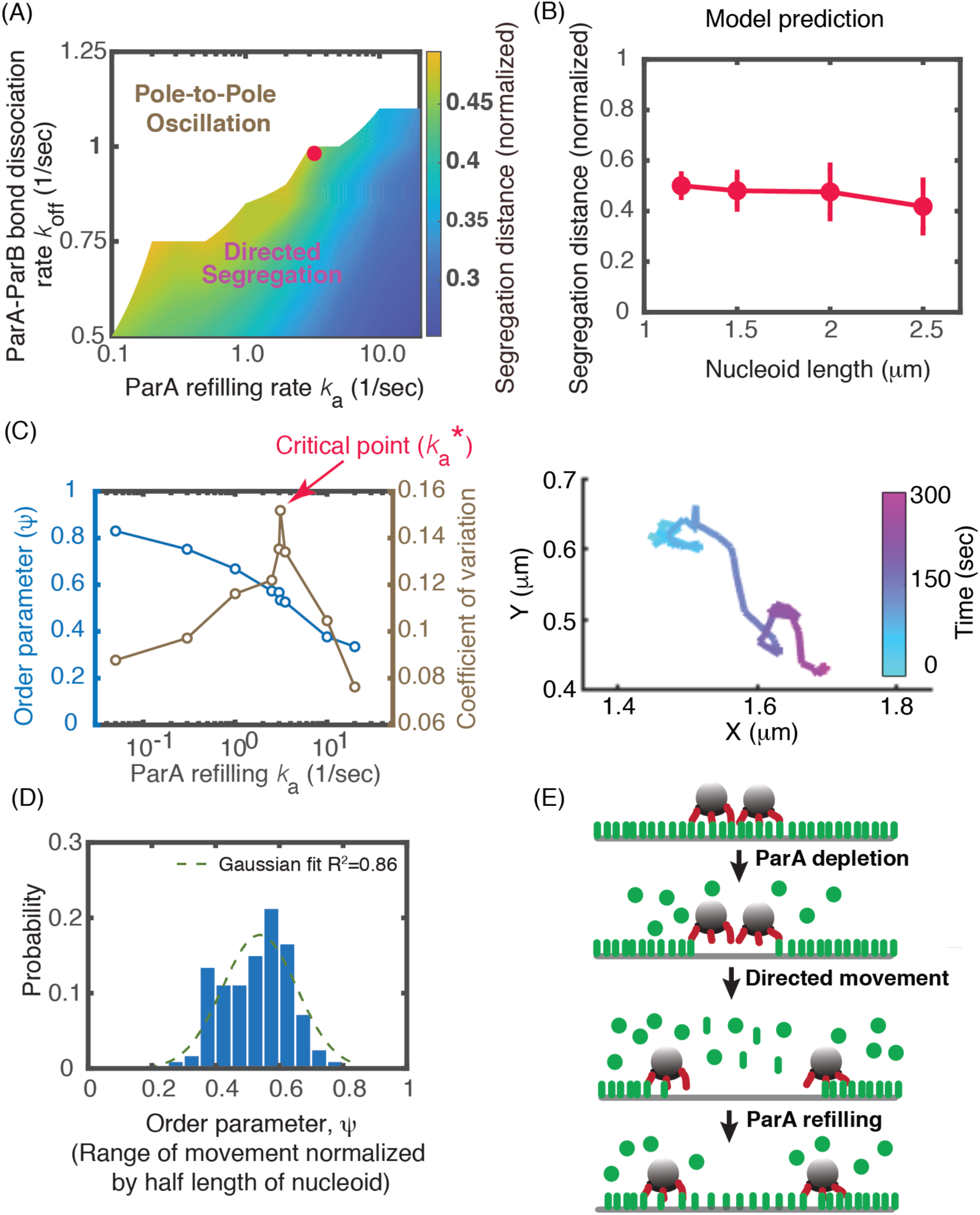
Predicted criticality of ParA-mediated PC partition. (A) Predicted phase diagram of PC motility controlled by (k_a_, k_off_). For each point in the phase diagram, we ran stochastic simulations for ≥ 36 trajectories of 10 min-dynamical evolution of the system, starting from the same initial condition and parameter set. The segregation distance reports the average value of ≥ 36 trajectories at the end of the simulation. Here, the ParA concentration is kept constant as ∼ 3500 molecules per micron of nucleoid length and other parameters are kept fixed (see Table S1 for details). (B) Segregation distance adapts to 1/2 of the nucleoid length at the critical point in the parameter space. (C) Predicted characteristics of critical point. Left: Order parameter (*i.e.*, the excursion distance of PC normalized by the half length of the nucleoid) increases continuously as the ParA refilling rate decreases, whereas the variance of the order parameter peaks around the critical point 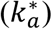. Note that the order parameter evolves similarly as a function of the ParA-ParB dissociation rate, k_off_. Right: Corresponding representative simulation trajectory of PC excursion with the parameter set around this critical point. (D) Predicted statistical distribution of order parameter ψ near the critical point (n=128). (E) Schematic illustration of the physical nature of near-critical-point partitioning.

Exploiting agent-based stochastic simulations of our model allows us to calculate the phase diagram of PC segregation motility with an elongating nucleoid (Fig. 2A). It is characterized by two key control parameters – the ParA-ParB bond dissociation rate, *k*_off_, and the ParA-nucleoid binding rate, *k*_a_. Our calculation suggested that the ParA gradient-based Brownian-ratcheting could allow the replicated PCs to undergo directed segregation and adapt their segregation distance at ∼ 0.5 of the increasing nucleoid length (Fig. 2B). This ensures the two PCs to always end up in the different cell halves, maximizing the partition fidelity. Importantly, this partition fidelity requires the ParAB*S* system to operate in a very special parameter regime (*e.g.*, the “•” denoted in Figure 2A), which represents the transition of PC motility from directed segregation to pole-to-pole oscillation. At/near this particular parameter set, the PCs are predicted to still undergo directed motility but with extensive excursions (Fig. 2C), rather than positioning around a specific landmark (Fig. 2 – Figure supplement 1A) nor oscillating perpetually all over the place (Fig. 2 – Figure supplement 1B). Importantly, the statistical distribution of the order parameter ψ (*i.e.*, the excursion distance normalized by the half length of the nucleoid) is very broad and highly non-Gaussian (Fig. 2D), which are distinctive features of critical dynamics (Goldenfeld 1992).

To test the prediction, we conducted time-lapse epifluorescence microscopy experiments using the well-established F-plasmid partition system in *E. coli* (Bouet and Funnell 2019; Guilhas et al. 2020). The plasmid F is present at ∼2 copies per chromosome per cell (Frame and Bishop 1971). We used functional fluorescent fusion proteins ParB_F_-mTq2 (Guilhas et al. 2020) and HU-mCherry to label the mini-F plasmid and the nucleoid, respectively (Fig. 3A). *E. coli* cells were grown in two different conditions giving raise to two generation times of 45 and 242 min with an average mini-F copy-number per cell of 3.6 and 2.8, respectively (see Table 1 in MATERIALS and METHODS). In these two conditions, we found that cells displayed 3.1 and 2.2 fluorescent foci per cell, respectively, indicating that during most of the cell cycle, plasmids are not clustered.

**TABLE 1.**
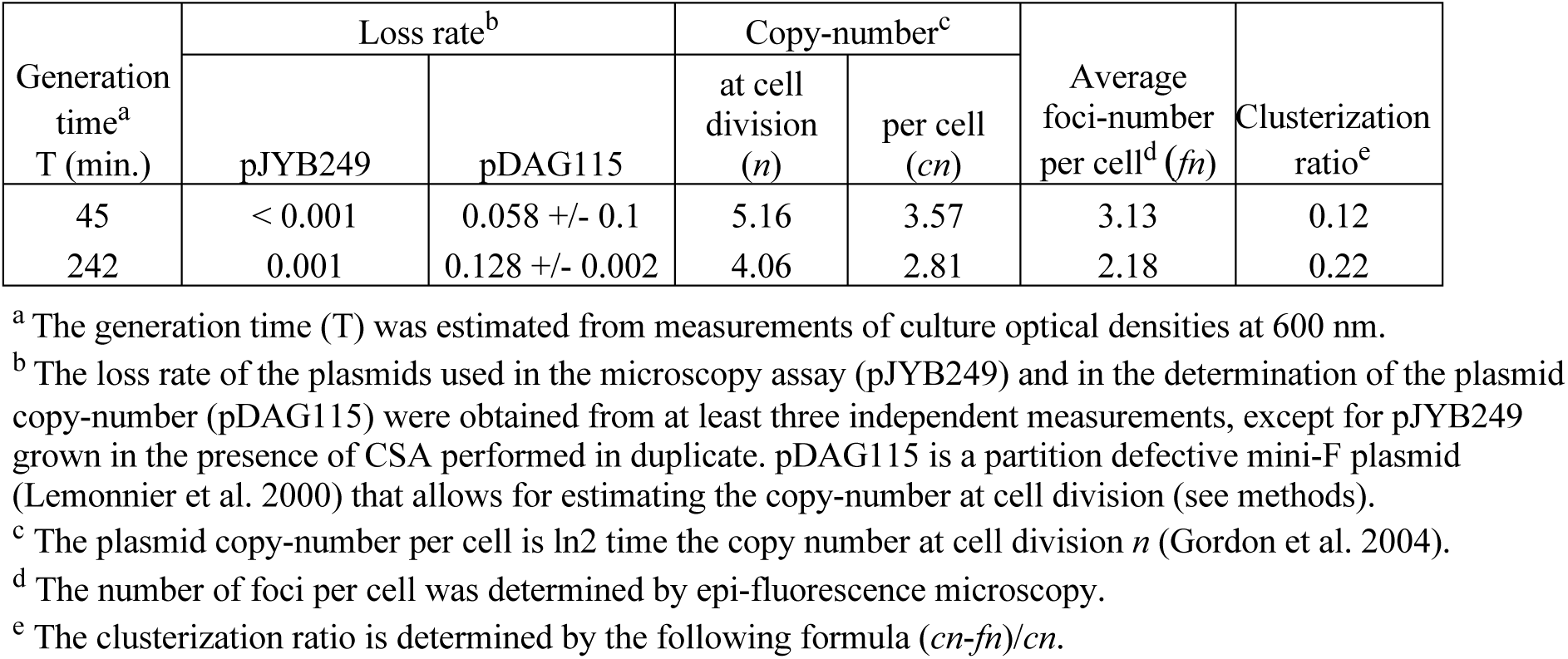
The level of plasmid F clusterization is low in fast and slow *E. coli* growing conditions. The clusterization level is estimated by the difference between the average number of foci and the average number of plasmids per cell.

| Generation time<sup>a</sup><br>T (min.) | Loss rate<sup>b</sup> | | Copy-number<sup>c</sup> | | Average foci-number per cell<sup>d</sup> ($\bar{f}_n$) | Clusterization ratio<sup>e</sup> |
| --- | --- | --- | --- | --- | --- | --- |
| | pJYB249 | pDAG115 | at cell division ($n$) | per cell ($cn$) | | |
| 45 | < 0.001 | 0.058 +/- 0.1 | 5.16 | 3.57 | 3.13 | 0.12 |
| 242 | 0.001 | 0.128 +/- 0.002 | 4.06 | 2.81 | 2.18 | 0.22 |

<sup>a</sup> The generation time (T) was estimated from measurements of culture optical densities at 600 nm.
<sup>b</sup> The loss rate of the plasmids used in the microscopy assay (pJYB249) and in the determination of the plasmid copy-number (pDAG115) were obtained from at least three independent measurements, except for pJYB249 grown in the presence of CSA performed in duplicate. pDAG115 is a partition defective mini-F plasmid (Lemonnier et al. 2000) that allows for estimating the copy-number at cell division (see methods).
<sup>c</sup> The plasmid copy-number per cell is $\ln 2$ time the copy number at cell division $n$ (Gordon et al. 2004).
<sup>d</sup> The number of foci per cell was determined by epi-fluorescence microscopy.
<sup>e</sup> The clusterization ratio is determined by the following formula $(cn - \bar{f}_n)/cn$ .

**Figure 3.**
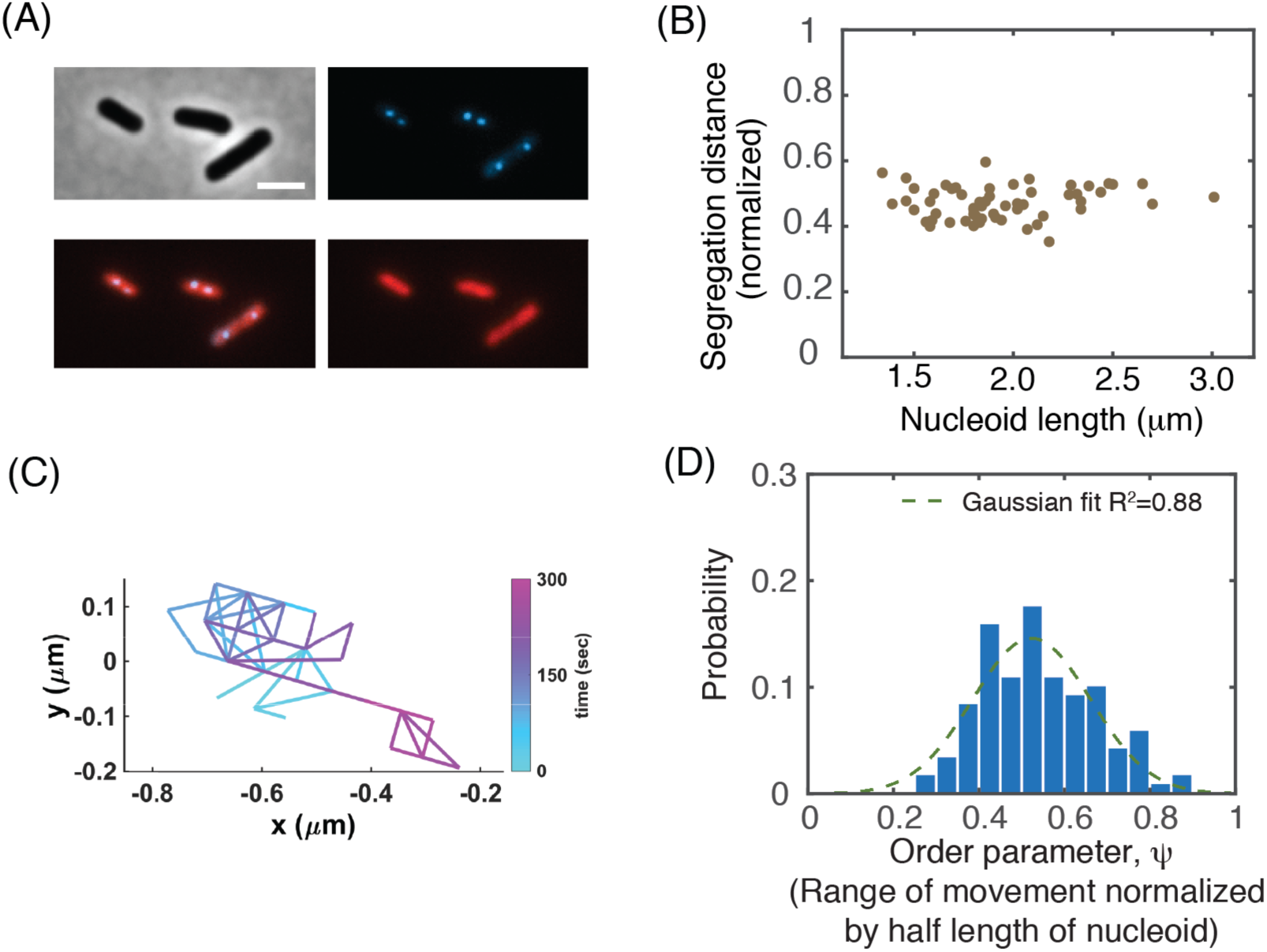
Critical-point-operation of ParA-mediated PC partition. (A) Experimental setup of two-color live-cell imaging of F-plasmids in wild-type *E. coli*. Cells are observed in phase contrast (top left) and in the blue (top right) and red (bottom right) channels for fluorescence microcopy to observe ParB_F_-mTq2 or ParA_F_-mVenus, respectively. Overlay of blue and red channel (bottom left); Scale bar (2 µm). (B) Experimental data demonstrate that PC segregation distance adapts to 1/2 of the nucleoid lengths (n=58). Data from cells grown at 30°C in MGly with or without casamino acids were represented on the same graph since they display the same trend. (C) A representative saltatory trajectory of PC excursion. (D) Non-Gaussian distribution of order parameter ψ (*i.e.*, the PC excursion range normalized by the half length of the nucleoid) (n=120).

Then, we measured the nucleoid size along the long cell axis and the distance between two PCs in two-foci cells. Our data showed that the PC segregation distance adapts to 0.5 of nucleoid length independently of the growth condition as the nucleoid increased from 1.2 micron to 3.0 microns (Fig. 3B). Critically, the PCs indeed undergo extensive excursion (Fig. 3C) and display a broad distribution (Fig. 3D). Further, we used excess kurtosis value, a common measure in statistics, to gauge how far the distribution deviates from a Gaussian distribution. The excess kurtosis value is 0 for the perfect Gaussian and −1.2 for the uniform distributions; in comparison, it is −0.34 for our experimental data and −0.68 for the model result (Fig. 2D and Fig. 3D). This indicates that these distributions deviate from a Gaussian distribution, which are telltale signs of criticality, confirming our prediction (Fig. 2D).

We interpret the physical nature of this near-critical-point PC partition as follows (Fig. 2E): Replicated PCs undergoes directed segregation because of their initial side-by-side arrangement. As they deplete the ParA underneath from the nucleoid, the local nucleoid-bound ParA concentration field becomes asymmetric in the “eyes” of each PC, which sets the directed movement. As the PCs move apart, each PC associates with a ParA-depletion zone, like “a sphere of influence”. It takes a while for the depleted ParAs to rebind the nucleoid, which eventually re-establish the symmetric ParA distribution surrounding each of the PCs, thus positioning the PCs. As such, the ParA-ParB bond dissociation confers the PC directed segregation, whereas ParA refilling event hinders it. The balance between these two activities defines the tipping point (*aka* the critical point in parameter space). At this tipping point, the PC’s ParA-depletion zones (or spheres of influence) overlap and span the entire nucleoid length. This allows the PCs to “feel” not only the presence of each other but the boundary of the elongating nucleoid.

Together, our results suggest that while the ParA-ParB interaction is local, operation of ParA gradient-based Brownian ratchet mechanism – near a critical point – can provide the PCs a global “view” that allows sensitive adaptation of their segregation distance to the increasing nucleoid length. That is, the near-critical-point operation allows ParA-mediated partition machinery to “measure” the cellular distance by molecular interactions.

### ParABS-mediated PC partition is robust against cell-to-cell ParA level variations

Given the sensitive nature of partitioning near a critical point, the next question is: How does the ParAB*S*-mediated partition manage to buffer against fluctuations inside cells, a central topic of control for any biological systems? To begin to characterize the cellular fluctuations relevant to ParAB*S*-mediated partition, we measured the intracellular ParA concentration by coupling ParA to m-Venus fluorescent peptide (ParA_F_-mVenus) in cells allowing to detect the nucleoid length and PC positioning (as above). The data showed that [ParA] varies from cell to cell over 10 folds in wildtype *E. coli* (Fig. 4A). Despite the large variations of [ParA], the PC partition still adapts the separation distance to ∼ 0.5 of various nucleoid lengths (Fig. 4A), ensuring the high partition fidelity observed with a plasmid loss rate ≤ 0.1% (see Table 1 in MATERIALS and METHODS).

**Figure 4.**
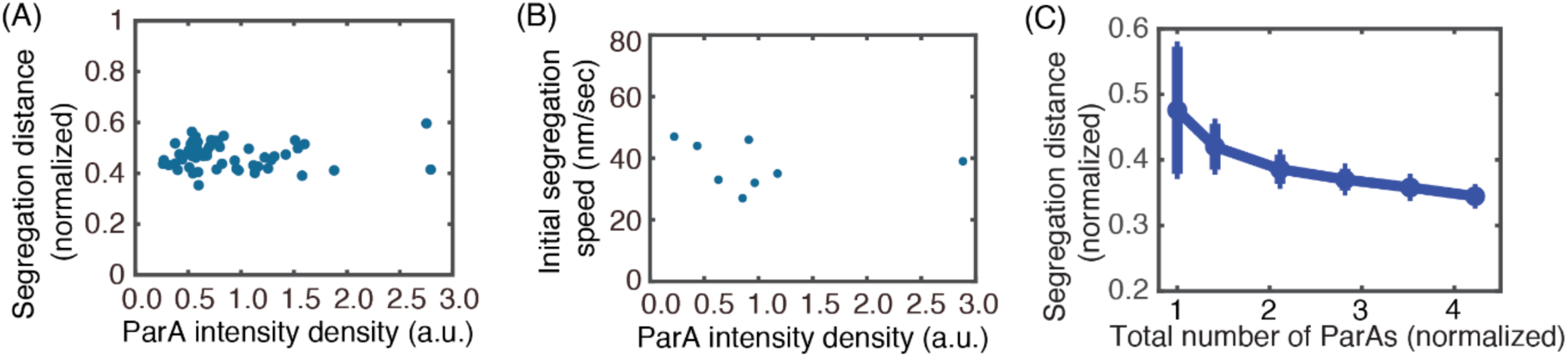
Robustness of PC partition against variations of ParA level. (A) Experimental data showing the PC segregation distance normalized by the nucleoid length is insensitive to the large cell-to-cell ParA level fluctuations (n=58). (B) Initial segregation speed is insensitive to the ParA level. (C) Current model cannot buffer the near-critical-point partition against ParA level fluctuations.

To understand the robustness measure adopted by the PC partition system, we reasoned that both the ParA-ParB dissociation rate, *k*_off_, and the ParA-nucleoid binding rate, *k*_a_ might change with the [ParA]. This way, the partition could still operate near the criticality when [ParA] varies; *i.e.*, rather than at a fixed point in the parameter space in Fig. 2A, the partition operates along the critical line between the states of pole-to-pole oscillation and directed segregation. To explore this possibility, we tried to measure the ParA_F_-ParB_F_ dissociation rate *k*_off_ by monitoring the PC foci segregation rates. According to our model, the PC segregation speed is proportional to the ParA_F_-ParB_F_ bond dissociation rate (Hu et al. 2017a). Our data show that the PC segregation speed, although varying somewhat, is insensitive to the [ParA] (Fig. 4B). Strikingly, this indicates that the *k*_off_ and hence, the *k*_a_ does not change much under these conditions. However, our current model showed that fixing the parameter set near the critical point, [ParA] variations would significantly perturb the adaptation of segregation distance to nucleoid length (Fig. 4C). This cannot be rescued by varying the values of *k*_off_ and *k*_a_ to the similar levels displayed in Fig. 4B. This discrepancy suggests partition robustness entails additional factor(s) and thus precipitates the question of what buffers the robustness of PC segregation against [ParA] variations.

### PC-mediated ParA localization underlies partition robustness against bulk [ParA] variations

Based on *in vitro* and *in vivo* data suggesting that ParA could bind to the plasmid through its interaction with ParB and ns-DNA (Bouet et al. 2007; Hatano and Niki 2010; Vecchiarelli, Hwang, and Mizuuchi 2013), our leading hypothesis is that the intracellular ParA could localize around the PC and create a local environment that buffers the partitioning against the bulk [ParA] variation (Fig. 5A). To test the hypothesis, we first extended the *in vivo* model to incorporate this PC-mediated ParA localization effect. We assumed the cytosolic ParA to have a high binding affinity to PC but with a transient lifetime before turning over into cytoplasm (Fig. 5A), characterized by the *k*_a, plasmid_ and *k*_d, plasmid_ rates. Due to the finite size of the PC, the model imposed an upper limit in the number of ParA molecules ∼ 100’s that simultaneously localize to the PC. This saturation level is based on the *in vitro* measurements (Hwang et al. 2013; Vecchiarelli, Hwang, and Mizuuchi 2013). We further assumed that right after releasing from the PC, the ParA has a reduced propensity to bind to the nucleoid. This last assumption is based on the observation that the interaction with ParB not only speeds up the dissociation of ParA from nucleoid but inhibits the ParA-nucleoid binding (Vecchiarelli, Hwang, and Mizuuchi 2013). Below, we will present the typical model result, the essence of which persists in a broad range of model parameter space (Fig. 5 – Figure supplement 1).

**Figure 5.**
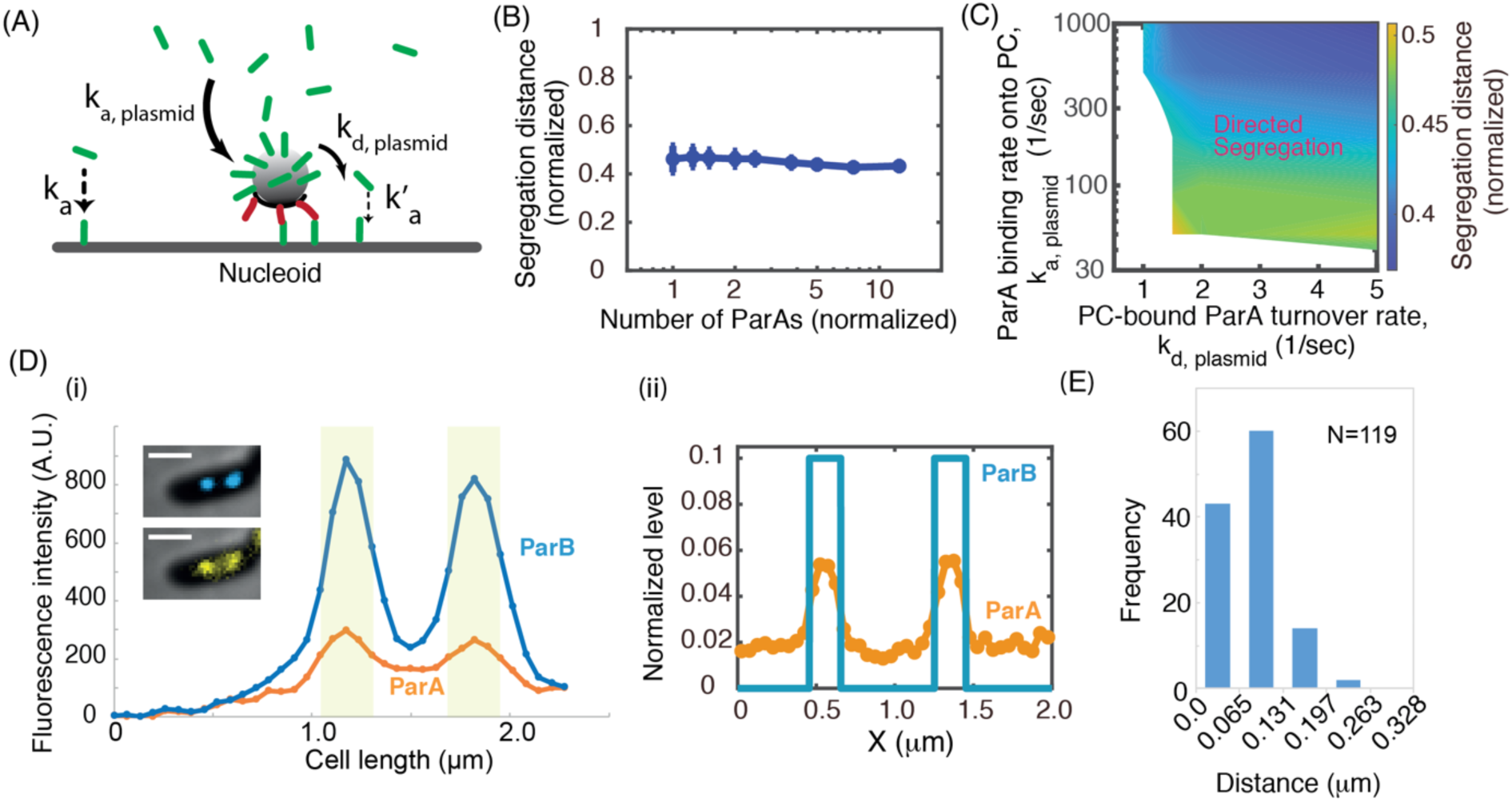
PC-localization of ParA explains the partition robustness. (A) Model scheme of PC-mediated ParA localization. (B) The amended model can ensure the fidelity of near-critical-point partition against the ParA level fluctuations. Note, the same parameter set here also simultaneously ensures the sensitive adaption of segregation distance to nucleoid lengths (see Fig. 5 – Figure supplement 2C). (C) Predicted phase diagram of the dependence of PC partition fidelity on the ParA localization effects. For each point in the phase diagram, we ran stochastic simulations for ≥ 36 trajectories of 10 min-dynamical evolution of the system, starting from the same initial condition and parameter set. The segregation distance reports the average value of ≥ 36 trajectories at the end of the simulation. (D) Representative spatial profiles of ParA and PC along the cell length. (i): Live-cell experimental result. Line scan analysis of the fluorescence intensities in arbitrary unit (A.U.) along cell length. Blue and orange lines correspond to the blue (ParB_F_-mTq2) and yellow (ParA_F_-mVenus) channels, respectively. The corresponding cell images is displayed in the graph with scale bar (1 μm). The grey area corresponds to 4 pixels (262 nm) around the PC peak. (ii): Model result. (E) Histogram of PC-ParA colocalization.

Equipped with this ParA localization effect, simulating this integrated mathematical model shows that it preserves the key features of criticality (Fig. 5 – Figure supplement 2A and B) and especially, the sensitive adaptation of PC segregation distance to the nucleoid lengths (Fig. 5 – Figure supplement 2C). More importantly, this PC-localization effects of ParA could simultaneously explain the robustness of partitioning evidenced in *E. coli.* (Fig. 4A). That is, the segregation of the replicated PCs by the half of the nucleoid length remains largely insensitive to the [ParA] variations (Figs. 5B and Figure supplement 2C). Such a buffering effect entails an appropriate ParA accumulation around the PC (Fig. 5C). When the on rate (*k*_a, plasmid_) is too slow, the PC not only depletes the ParA from underneath but cannot supply enough ParA to refill the depletion zone underneath so that the PC becomes diffusive (the lower portion of Fig. 5C). When the on rate is too fast and the off rate is too slow, the PC will accumulate too many ParAs so that the nucleoid-bound ParA will become very sparse, likewise favoring diffusive movement (the upper left corner of Fig. 5C). However, as the off rate increases while keeping the on rate very fast, the PC will funnel the cytoplasmic ParA to the local nucleoid at very high concentration. This significantly increases the overall ParA binding to nucleoid, immobilizing the PCs (the upper right corner of Fig. 5C). In these extreme limits, the partition system loses its robustness of adapting the PC segregation distance to the half of an elongating nucleoid (Fig. 5C). To ensure the PC partition fidelity, the ParA thus is expected to localize around the PCs with only several-fold accumulation at its peak concentration.

To begin to test this prediction, we resorted to live-cell imaging to discern whether and how ParA accumulates around the PC. To better resolve the sub-cellular pattern, we sought the experimental conditions that have a low amount of ParA without perturbing other key factors of the system. We took the advantage of our observation that in wildtype cells ParA distributes asymmetrically between the two daughter cells at cell division (Fig. 5 – Figure supplement 3A), which not only underlies the large cell-to-cell variation of [ParA] (Fig. 4A) but presents a natural testing ground of our model. We thus focused our analyses on the new-born cells that have inherited a low amount of ParA_F_-mVenus (Fig. 5 – Figure supplement 3A), which allows for a better detection of localized signals. In these cells with low intracellular amount of ParA, we also imaged the PC locations with ParB_F_-mTq2 that form intense foci. We measured the peak intensity along the cell length for both ParA and ParB by applying a line scan analysis. Although ParA-mVenus displays faint foci, they appeared very close to PCs; importantly, they did not result from cross-fluorescence imaging (Fig. 5 – Figure supplement 3B). Fig. 5D(i) presents a representative measurement and Fig. 5 – Figure supplement 3C provides more detailed analysis. Briefly, in the 58 cells analyzed, we observed that the vast majority of them (41 cells) display the same number of ParA and ParB foci, 14 cells display one more ParA focus than ParB, and 3 cells have one more ParB focus than ParA. However, we could not accurately measure the ParA intensity accumulation around PC because of (i) the cell to cell variation in [ParA] and (ii) the depletion of ParA provoked by ParB. Nevertheless, the observation of discrete ParA patches indicates that ParA accumulates several folds compare to the intracellular level in the close vicinity of PCs, consistent with our predictions (Fig. 5D(ii)). To further quantify the degree of co-localization of the ParA and ParB foci, we measured the distance between the peak intensities for each pair of ParA and ParB foci (n = 119; Fig. 5E). We found that for 98% of the pairs, the ParA and ParB peak intensities are within 2 pixels (131 nm). Given the 200-250 nm resolution of epifluorescence microscopy due to light diffraction limit, our data show that ParA and ParB foci are highly co-localized, providing strong support to the predicted PC-ParA co-localization. Importantly, regardless of the ParA levels (Fig. 4 and Fig. 5 – Figure supplement 3), the corresponding PC segregation distance always adapts to ∼ half of the nucleoid length (Fig. 3A) with a very low error rate of the partitioning (< 0.1%) (see Table 1 in MATERIALS and METHODS). Combining our model and experimental data, we suggest that PC-mediated ParA localization underlies the fidelity of PC partitioning against [ParA] variations.

## DISCUSSION

In this paper, we provided direct evidences – with a mechanistic underpinning – that the partitioning of low-copy-number plasmids operates near a critical point. The near-critical-point operation allows the partition machinery to gauge the size of the entire nucleoid and accordingly, adapt the plasmid segregation distance to the half-length of the elongating nucleoid (Fig. 3). Segregating by half of the nucleoid length renders that each cell halves always inherit at least one PC, ensuring the partition fidelity, which is also observed in other ParABS systems (*e.g.*, (Ietswaart et al. 2014)). We further provided the data suggesting that the PC localizes cytoplasmic ParA to its neighborhood. This spatial control creates a local environment that buffers the near-critical-point partition against the fluctuations in bulk [ParA] (Fig. 5), which allows the cell to manage the dichotomy of sensitive adaption and robust execution of low-copy-number plasmid partitioning. This way, each PC defines its own “sphere of power” that allows the PC to “self-drive” by generating the path ahead and erasing the trail behind.

The current model predicts ParA-PC localization effects by only focusing on the simplest biochemical scheme. Nonetheless, we indeed observed the ParA-PC localization when the ParA level is low in wild-type cells (Fig. 5D and Fig. 5E). Given that 1) PC is much smaller in size than the nucleoid and 2) it competes with the nucleoid to bind the same pool of cytoplasmic ParA, this observation indicates that ParA-PC binding affinity must be much higher than its nucleoid-counterpart. Since ParA binds to ns-DNA, without preference for plasmid over chromosome DNA, its co-localization with PC is expected to arise from its interactions with the PC-bound ParB. This notion is consistent with the extensive experimental evidence that the PC-bound ParB interacts with the cytoplasmic ParA (Bouet and Funnell 1999; Figge, Easter, and Gober 2003; Osorio-Valeriano et al. 2019). We thus expect ParA to localize around the PC when its level increases in wild-type cells, although it would be difficult to experimentally discern its localization pattern with a high background. Importantly, according to our model (Fig. 5B and Fig. 5C), this PC-localization of ParA ensures the fidelity of PC partition regardless of the ParA level variations that naturally occur in wildtype cells (Table 1 and Fig. 4A).

The direct testing of how the PC-localized ParA drives the partition fidelity of the *parS*-carrying DNA is not simple. Point mutations that disrupt ParA–ParB interactions and compromise ParA ATPase activity were identified, *e.g.*, ParA_F_-K120Q and ParA_F_-K120R (Libante, Thion, and Lane 2001). These mutations that drastically changed the spatial profile of ParA_F_ from foci (Fig. 5D) to uniform distribution along the nucleoid (Hatano, Yamaichi, and Niki 2007) were reported to increase the loss rate of the F-plasmids by 400-800 folds (Libante, Thion, and Lane 2001). These observations would be consistent with the model proposal that decreasing the PC-localization of ParA compromises the partition fidelity (*i.e.*, the lower left corner of Fig. 5C). We caution, however, that the point mutations render the PC to lose contact with the nucleoid and, hence, the active partition after all (Le Gall et al. 2016). It is possible that localizing to PC is integral of ParA’s normal activities. Thus, identifying a ParA variant that is specifically perturbed in its PC-localization but not in other activities essential to partition is not easily accessible. Such a study is also be complicated with the transient interactions between ParA and ParB that both exist in different biochemical states (Vecchiarelli et al. 2010; Vecchiarelli, Hwang, and Mizuuchi 2013; Ah-Seng et al. 2009). The recent findings that ParB binds CTP (Soh et al. 2019; Osorio-Valeriano et al. 2019; Jalal, Tran, and Le 2020) and that *parS*-mediated CTP hydrolysis stimulates ParA interaction (Osorio-Valeriano et al. 2019; Jalal, Tran, and Le 2020) could open interesting molecular clues to control this interaction.

Moreover, our model describes the PC partitioning along the nucleoid surface as a 2D problem. Recent experiments, however, suggest that the PC may move inside the nucleoid (Le Gall et al. 2016). Let us consider the simplest 3D case first, followed by more complex scenarios. The simplest 3D scenario is that the interactions between the PC-bound ParB and nucleoid-bound ParA are still along the surface of the PC and the nucleoid should have the void space that allows the PC to move through. In this scenario, the PC moving through the nucleoid is like a sphere moving through a hollow cylinder. Now unfolding the hollow cylinder will render a flat substrate, akin to our current 2D model. Within this simplest 3D case, the 3D effects could alter several key parameters of the current model. First, trapping inside the nucleoid likely slows down the diffusion of the PC. Second, the number of ParA-ParB bonds in 3D might be different from its 2D-counterpart, as some ParB buried within the partition complex may not be available for bond formation with the nucleoid-bound ParA. Third, the nucleoid DNA density, instead of uniform, is reported to have high and low density-regions (Le Gall et al. 2016). In this regard, our phase diagram studies show that the essence of our conclusion – *i.e.*, the nature of critical-point-operation of PC partition and the effect of PC-localization of ParA on the robustness against [ParA] variations – is largely preserved against variations of these relevant model parameters (Fig. 5 – Figure supplement 4).

We note that a more realistic 3D model may contain many factors that are not well-characterized. For instance, we do not know the interior landscape of the nucleoid, nor whether and how the nucleoid DNA changes its conformation to accommodate the PC movement. Additionally, instead of acting as impenetrable structures, PC and nucleoid could be amorphous so that ParA and ParB are able to freely enter and exit, offering a more complex interaction network that remains to be explored. Given these uncertainties, we will leave the 3D model to our future study.

Furthermore, this PC-mediated ParA localization effect has its own limitation in ensuring the partition fidelity (Fig. 5 – Figure supplement 1). For instance, when the nucleoid becomes too long (Fig. 5 – Figure supplement 1D), the current model would predict the partition machinery to lose its ability of adapting the segregation distance to half of the nucleoid length. Interestingly, as the nucleoid gets longer, the low-copy plasmids are reported to replicate accordingly to keep the plasmid-chromosome ratio constant. The latter events increase the number of PC foci. This way, each PC focus “command” a unitary length of nucleoid as its power of sphere, resulting in the equi-distant pattern (Ebersbach et al. 2006; Sengupta et al. 2010): the two PCs on the same nucleoid segregate by 1/2 of the nucleoid length, whereas the three PCs will segregate by ∼ 1/3 of that nucleoid length, and so on. There emerges a size-scaling between the PC inter-distant and the nucleoid length, as a function of PC foci numbers. While the notations of size-scaling and size-control has indicated in biological systems (Crowder et al. 2015; West and Brown 2005; Chan and Marshall 2012; Willis and Huang 2017; Si et al. 2019), our work here provides a functional perspective, in a way similar to spindle-cell size scaling (Chen and Liu 2016).

Lastly, because low-copy-number plasmids provide selective advantages for bacterial survival, connecting the physical mechanism with the fidelity of DNA partition could allow us to understand how evolution might shape the near-critical-point behavior of a biological process to maximize its function. Interestingly, while widely conserved in both genome segregation (*i.e.*, plasmids and chromosomes) and subcellular organelle trafficking in bacteria (MacCready et al. 2018; Vecchiarelli, Mizuuchi, and Funnell 2012), the ParAB*S*-mediated partitioning displays distinct spatial-temporal features in these different systems (Fogel and Waldor 2006; Lim et al. 2014; Marston and Errington 1999; Schofield, Lim, and Jacobs-Wagner 2010). With these diverse spatial-temporal dynamics, our work provides a starting point to shed light on how the near-critical-point operation of the same machinery adapts to different systems with different sizes and geometries. In a broader context, the principles of spatial controls over the near-critical-point operation provide a possible solution to a fundamental question of cell biology: How do cells faithfully measure cellular-scale distances by only using molecular-scale interactions? We will investigate along this direction in the near future.

## MATERIALS AND METHODS

### Quantitative model formulation and simulation details

To quantitatively elucidate the proposed mechanism, we numerically compute our model with the parameters capturing *in vivo* conditions (Fig. 1 – Table supplement 1 in Appendix). While increasing over time, the bulk-part number of ParA molecules in the nominal case is set to be ∼ several thousands in the model, in accordance to the measurements (Adachi, Hori, and Hiraga 2006; Bouet et al. 2005). The nucleoid-bound ParA·ATPs were initially in chemical equilibrium with their cytosolic counterparts and were randomly distributed on the nucleoid lattice with 5 nm spacing. ParBs were permanently distributed with a uniform density of ∼ 0.013 ParB dimer/nm^2^ over the PC, quantitatively reflecting the measured high propensity of ParB to spreading around the *parS* site on the plasmid (Sanchez et al. 2015; Graham et al. 2014; Rodionov, Łobocka, and Yarmolinsky 1999; Lynch and Wang 1995). We model each ParA-ParB bond as an elastic spring. The vertical distance between the nucleoid and the PC was fixed at the equilibrium length of ParA-ParB bond (*L*_e_). The stochastic reactions involving PC-bound ParB, nucleoid-bound and cytosolic ParAs are simulated with the kinetic Monte Carlo scheme according to the reaction scheme (Fig. 1B).

The simulation workflow is as follows. At each simulation time step, each ParB can interact with available ParA·ATP within a distance *L*_*a*_, and bind only one ParA at a time with a rate of *k*_on_ (Step 1 in Fig. 1C). The probability of binding is proportional to 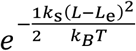 for *L*_a_ > *L* > *L*_e_; otherwise, it is zero. *K*_*s*_ is the spring constant of the bond, *L* denotes the separation between ParB and ParA·ATP. If this bond forms, *L* is the instantaneous bond length, 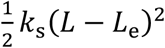 represents the associated elastic energy penalty. Importantly, given the model parameters, this energy penalty is less than the thermal energy, *k*_*B*_*T*. Consequently, thermal energy is sufficient to pre-stretch the newly formed bond, which in turn provides an elastic force *f* = *k*_*s*_(*L*– *L*_e_) (Step 2 in Fig. 1C). In the simulation, we vector-sum the elastic forces from all the ParA-ParB bonds over the PC. This net force together with the PC diffusion drives PC motion for one simulation step (Step 3 in Fig. 1C), following the Langevin-like dynamics: 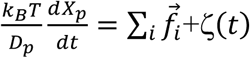. Here, *X*_p_ is the centroid position of the PC, *D*_p_ is the diffusion constant of the PC, 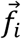 is the elastic force from the *i*th ParA-ParB bond on the PC, and ζ(*t*) represents random force resulting from thermal motion of the solvent molecules with 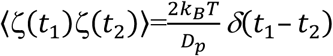 and *δ*(*t*_1_– *t*_2_) is the Dirac delta function.

In the next time step, the lengths and orientations of the ParA-ParB bonds are updated by the PC motion, from which the dissociation rates of the existing ParA-ParB bonds are calculated: when the bond extension exceeds the maximum, *X*_*C*_ (*i.e.*, (*L* – *L*_*e*_) > *X*_*C*_), the bond breaks instantaneously; otherwise, the dissociation rate is *k*_off_. This dissociation reaction is next implemented in the stochastic simulation. The resulting ParA^D^ will be released into cytoplasm and convert at a slow rate of *k*_*n*_ (*k*_*n*_ *≪ k*_*off*_) to ParA·ATP. Additionally, new ParA·ATPs are generated in cytosol concurrently with the elongation of nucleoid and cytoplasmic domains at the rate of *k*_nl_. These ParA·ATPs can bind to the nucleoid with a rate of *k*_*a*_.

Meanwhile, PC movement from the previous time step permits PC-bound ParBs to explore new territory and form bonds with available ParA·ATPs, and vacancies on the nucleoid can be re-filled by ParA·ATP rebinding from the cytosol or diffusing from adjacent sites on the nucleoid. These ParA·ATPs can establish new bonds with ParB if the PC is nearby. We then update the net force from all the ParA-ParB bonds, including changes in existing bonds and newly formed bonds. The movement of the cargo is then calculated as in the previous time step. We repeat these steps throughout the simulation over time.

### Bacterial strains and plasmids

*E. coli* K-12 strains are derivatives of DLT1215 (Bouet, Bouvier, and Lane 2006) and transformed with the plasmids pJYB240 (Guilhas et al. 2020), pJYB243 (Sanchez et al. 2015) or pJYB249 (Guilhas et al. 2020). Cultures were grown at 37°C with aeration in LB (Miller 1972) containing thymine (10 µg.ml-1) and antibiotics as appropriate: chloramphenicol (10 µg.ml^−1^); kanamycin (50 µg.ml^−1^). For microscopy and plasmid stability assays, cultures were grown at 30°C with aeration in MGlyC (M9 minimal medium supplemented with 0.4 % glycerol, 1 mM MgSO_4_, 0.1 mM CaCl_2_, 1 µg.ml^−1^ thiamine, 20 µg.ml^−1^ leucine and 40 µg.ml^−1^ thymine) supplemented or not with 0.2 % casamino acids. The generation times in MGly with or without 0.2 % casamino acids are 45 or 242 min, respectively.

### Plasmid stability assays for plasmid copy-number determination

Experiments were started from colonies of *E. coli* cells carrying the plasmids under test. Overnight cultures in M9Gly, with or without casamino-acids, containing chloramphenicol were diluted 250-fold into the same medium and grown to A_600_ = 0.25. Samples were then diluted serially into fresh medium without chloramphenicol and were processed as described previously (Lemonnier et al. 2000). To determine the fraction of cells that retained the plasmid, samples were taken at the beginning and after 5, 10, 20 and 30 generations or after 25 for growth in the absence or presence of casamino-acids, respectively. The loss frequency (*f*) per generation is calculated using the following formula: *f* = 1 – (cell F^+^ / total cell)^1/*g*^, where *g* is the generation number, as previously described (Sanchez et al. 2015).

The plasmid copy-number at cell division (*n*) is calculated from the probability of having one plasmid-free cell at cell division as a function of the copy-number, P_0_ = 2^(1-*n*)^, from which we obtained the theoretical frequency of random loss per generation. The copy number per cell is ln2 time *n* (Gordon et al. 2004).

### Epifluorescence microscopy and analysis

Overnight cultures were inoculated into fresh media at a concentration permitting at least 10 generations of exponential growth and incubated at 30°C to an optical density (OD_600_) of ∼0.25. Samples (0.7 μl) were deposited to the surface of a layer of 1% agarose buffered in M9 solution, as described (Diaz, Rech, and Bouet 2015). The cells were imaged at 30°C using an Eclipse TI-E/B wide field epifluorescence microscope with a phase contrast objective (CFI Plan APO LBDA 100X oil NA1.45) and a Semrock filter YFP (Ex: 500BP24; DM: 520; Em: 542BP27) or FITC (Ex: 482BP35; DM: 506; Em: 536BP40). Images were taken using an Andor Neo SCC-02124 camera with illumination at 80% from a SpectraX source Led (Lumencor) and exposure times of 0.2-1 second. Nis Elements AR software (Nikon) was used for image capture and editing. Image analyses were performed using Image J plugins. The average foci number per cell was measured using the ‘Cell counter’ plugins. Tracking the PC and nucleoid length was done in the MicrobeJ plugin (Schindelin et al. 2012; Ducret, Quardokus, and Brun 2016). It involves finding the cells of interest and tuning the parameters in MicrobeJ to get the data. Specifically, for each identified cell, we recorded the length of the cell, integrated fluorescence of ParA, and the peak positions of ParB fluorescence intensity, which were then used to represent the positions of the partition complexes. We also recorded the fluorescence profile of the nucleoid along the long axis of the cell, from which the nucleoid length of the cell was derived using the full-width-half-height approach. For ParA-PC colocalization analysis, we used a self-developed (CBI) Python-based tool (Distance2MaxProfile: https://imaprocess.pythonanywhere.com/Analymage/detail_projet/20/).

## ACKNOWLEDGEMENTS

The authors thanks Drs. Anthony Vecchiarelli, Keir Neuman and Kiyoshi Mizuuchi for the inspiring critiques of the work, and Christian Rouvière (ImageAnalysis-CBI) for implementing a python-based analysis tool for colocalization measurements.

## APPENDIX

### SECTION I. MODEL PARAMETER TABLE

**Fig. 1 – Supplemental table 1.**
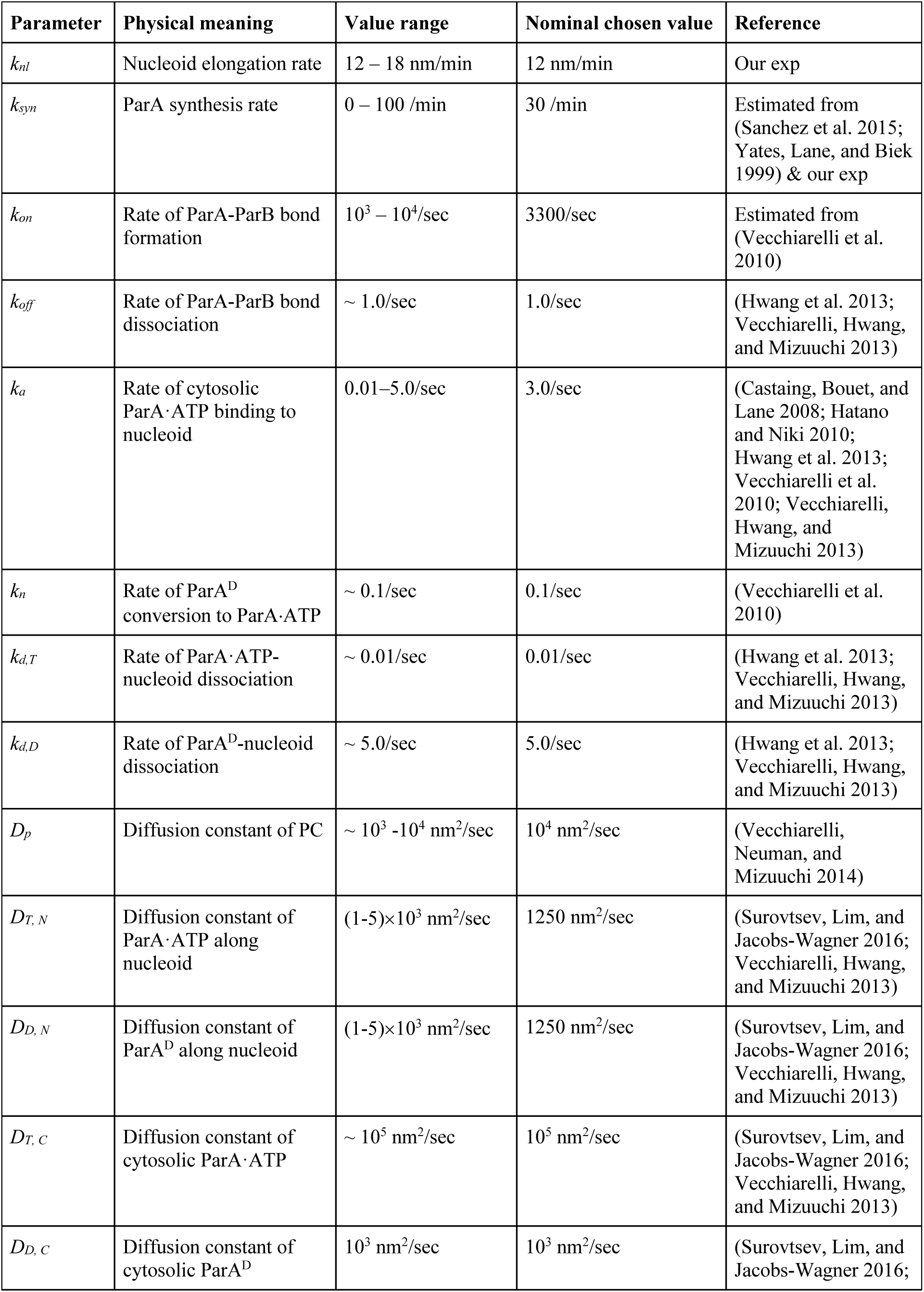
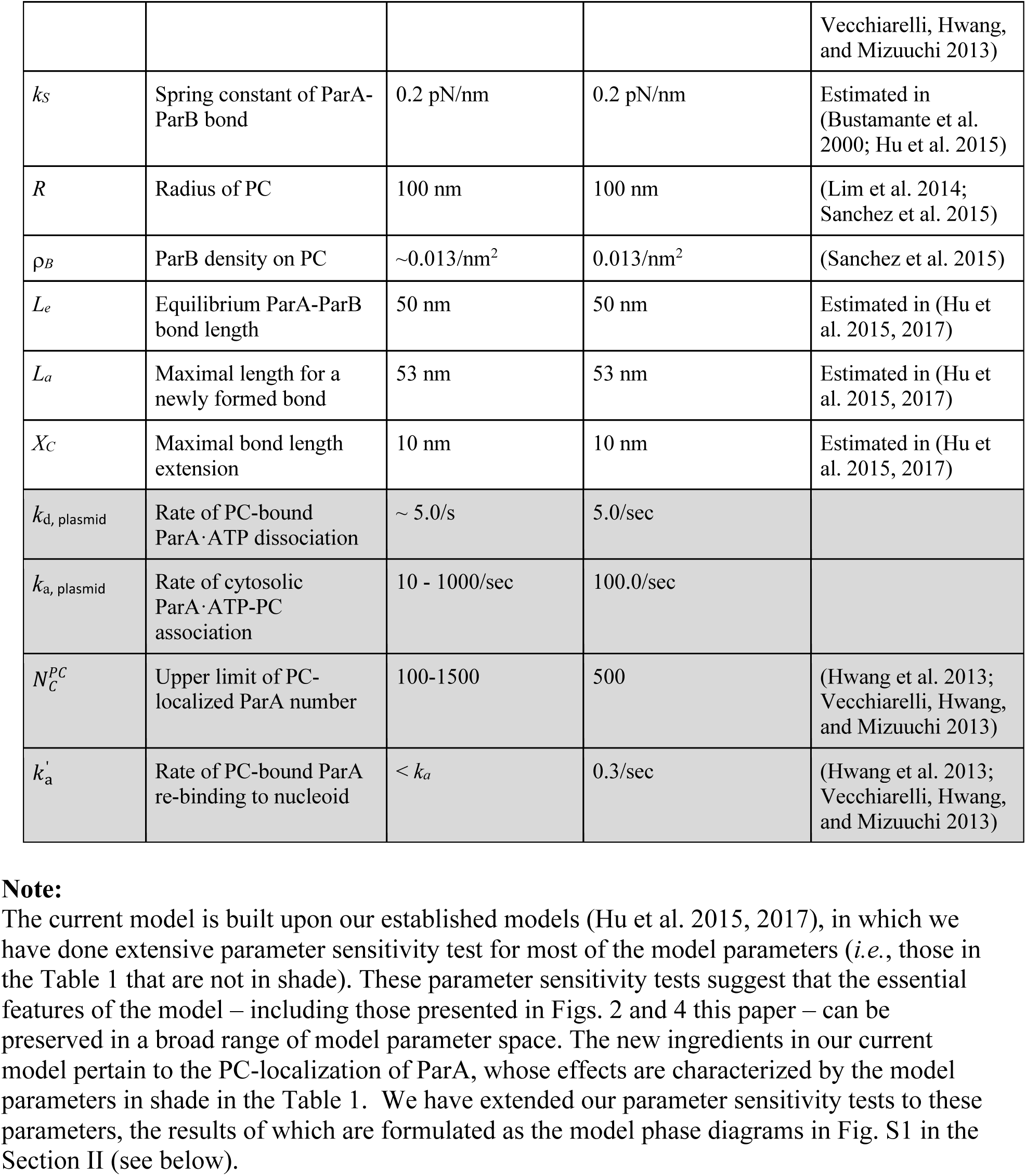

## SECTION II. SUPPLEMENTAL FIGURES

**Fig. 2 – Figure supplement 1.**
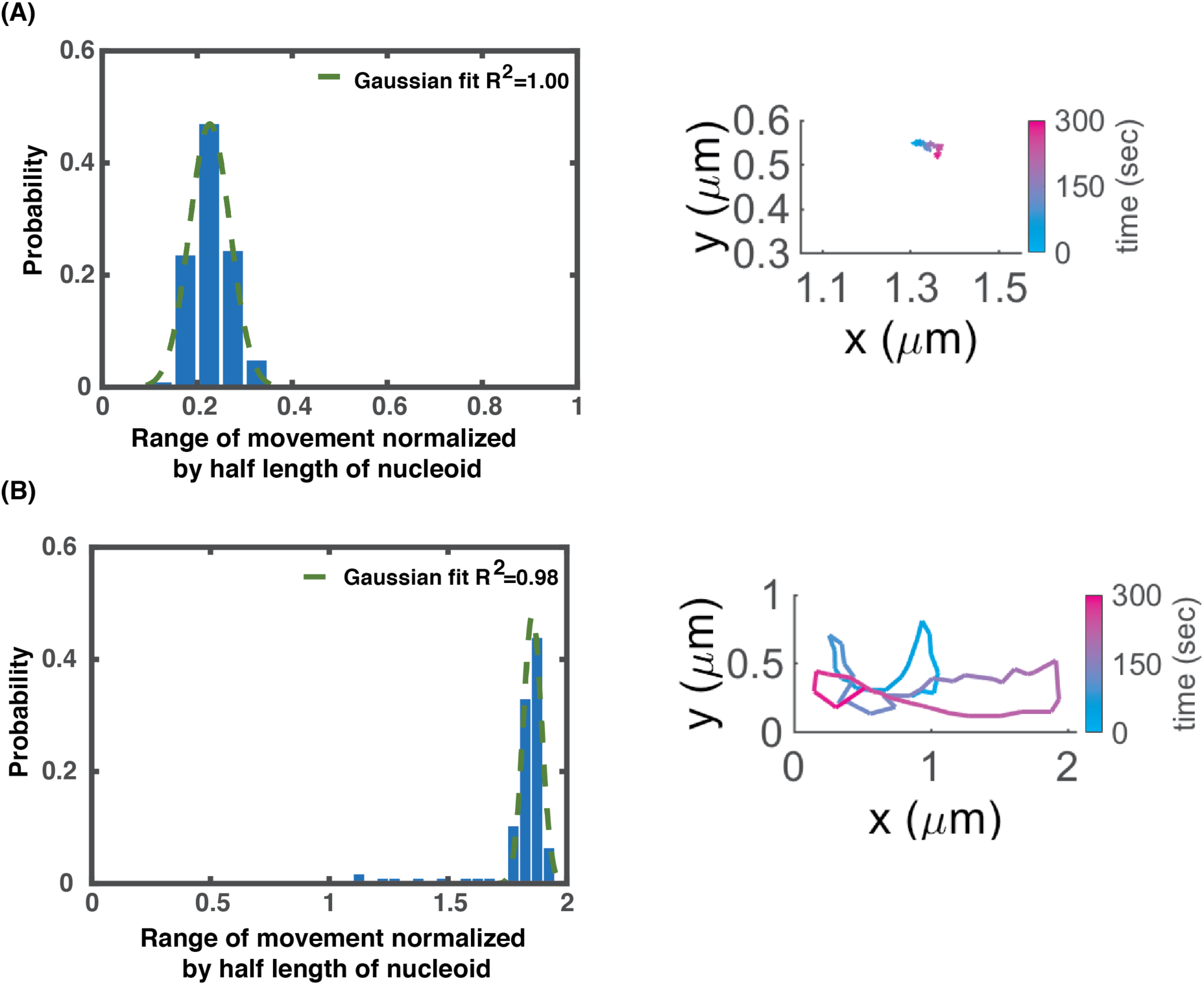
Statistical distributions of PC excursion distance (left) and representative simulation trajectories of PC movement (right) in the parameter regimes away from the critical point: (A) Near-static movement. (B) Pole-to-pole oscillation.

**Fig. 5 – Figure supplement 1.**
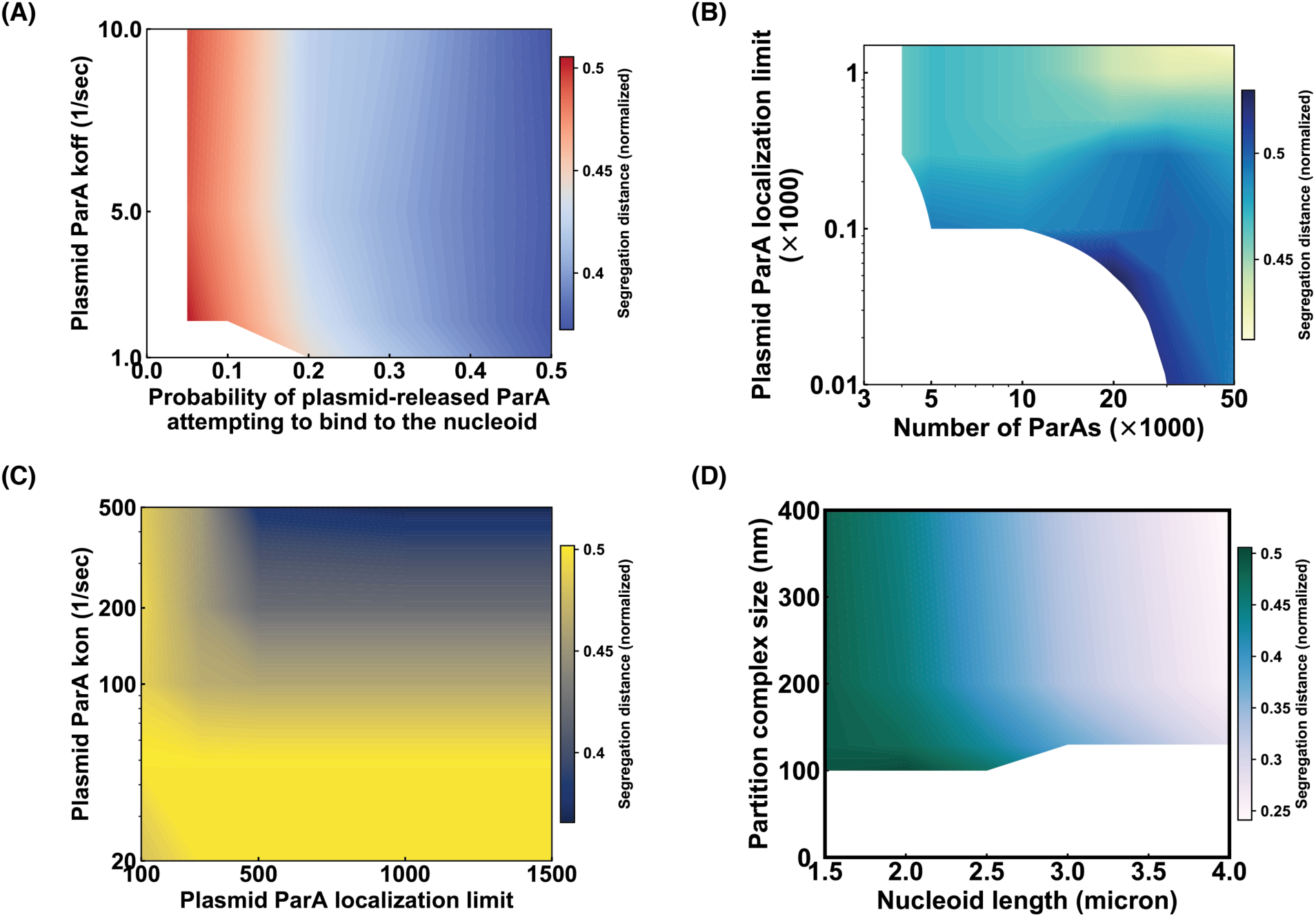
Model phase diagram studies of PC-ParA localization effects on PC partitioning. (A). ParA turnover rate from PC vs. percentage of PC-released ParA that binds to nucleoid. (B). Saturation level of PC-localized ParA vs. total ParA number. (C). ParA binding rate onto PC vs. Saturation level of PC-localized ParA. (D) PC size vs. nucleoid length. Here, the nucleoid length refers to the initial length of the nucleoid length in the simulation, which will elongate at the rate of 12 nm/min. For (A-D): We varied the two parameters in the respective phase diagrams, while keeping the rest of the model parameters fixed in accordance to the nominal chosen values in the Table 1. For each point in the phase diagram, we ran stochastic simulations for ≥ 36 trajectories of 10 min-dynamical evolution of the system, starting from the same initial condition and parameter set. The segregation distance reports the average value of ≥ 36 trajectories at the end of the simulation.

**Fig. 5 – Figure supplement 2.**
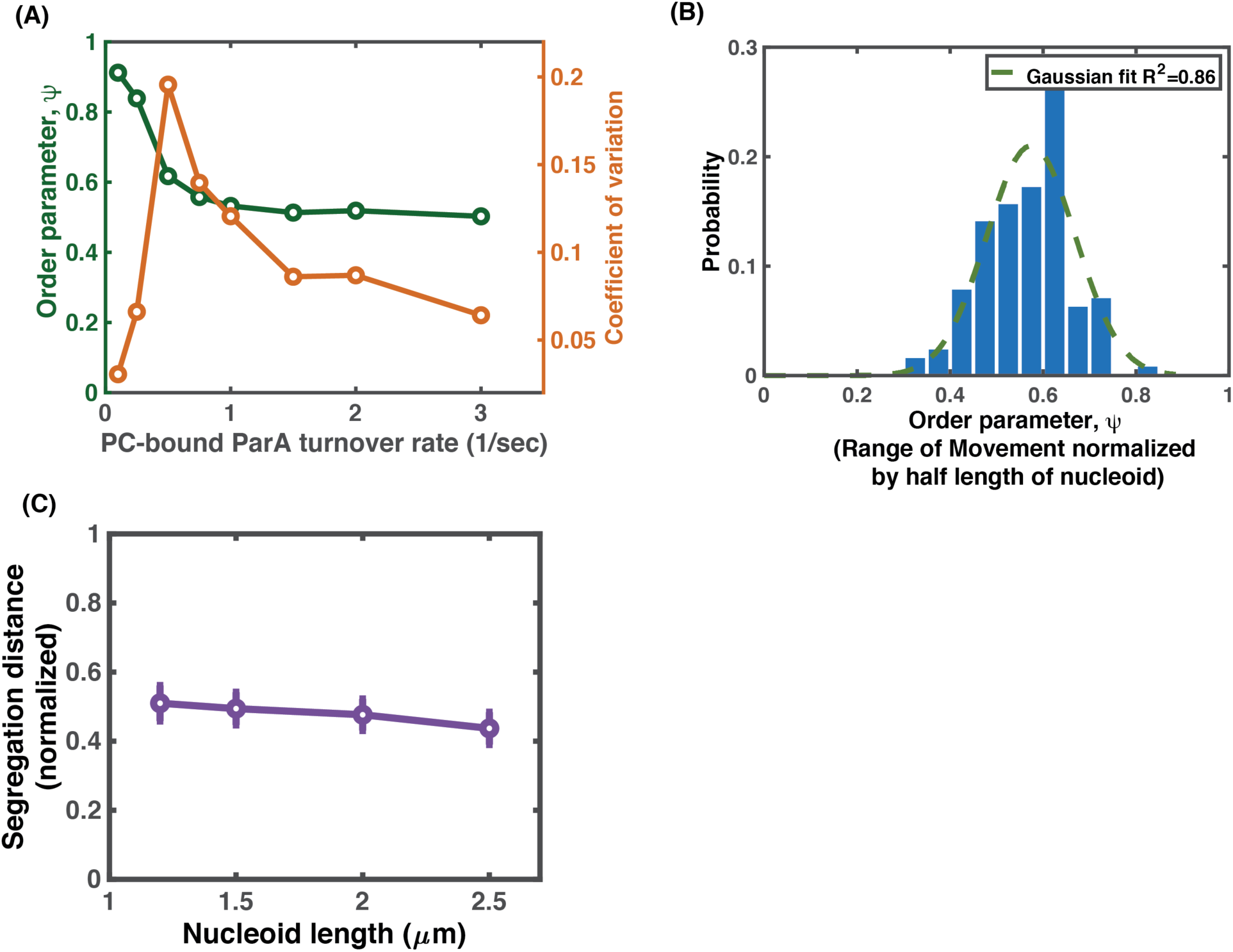
Full model with PC-localization of ParA preserves all the essence of near-critical-point partition as that in Figure 2. (A) Order parameter and its variation as functions of PC-bound ParA turnover rate. (B) Statistical distribution of the excursion distance near the critical point. (C) Segregation distance adapts to half of the nucleoid lengths near the critical point in the parameter space. The parameter set (*k*_a, plasmid_, *k*_off, plasmid_) used here is the same as that in Fig. 5B.

**Fig. 5 – Figure supplement 3.**
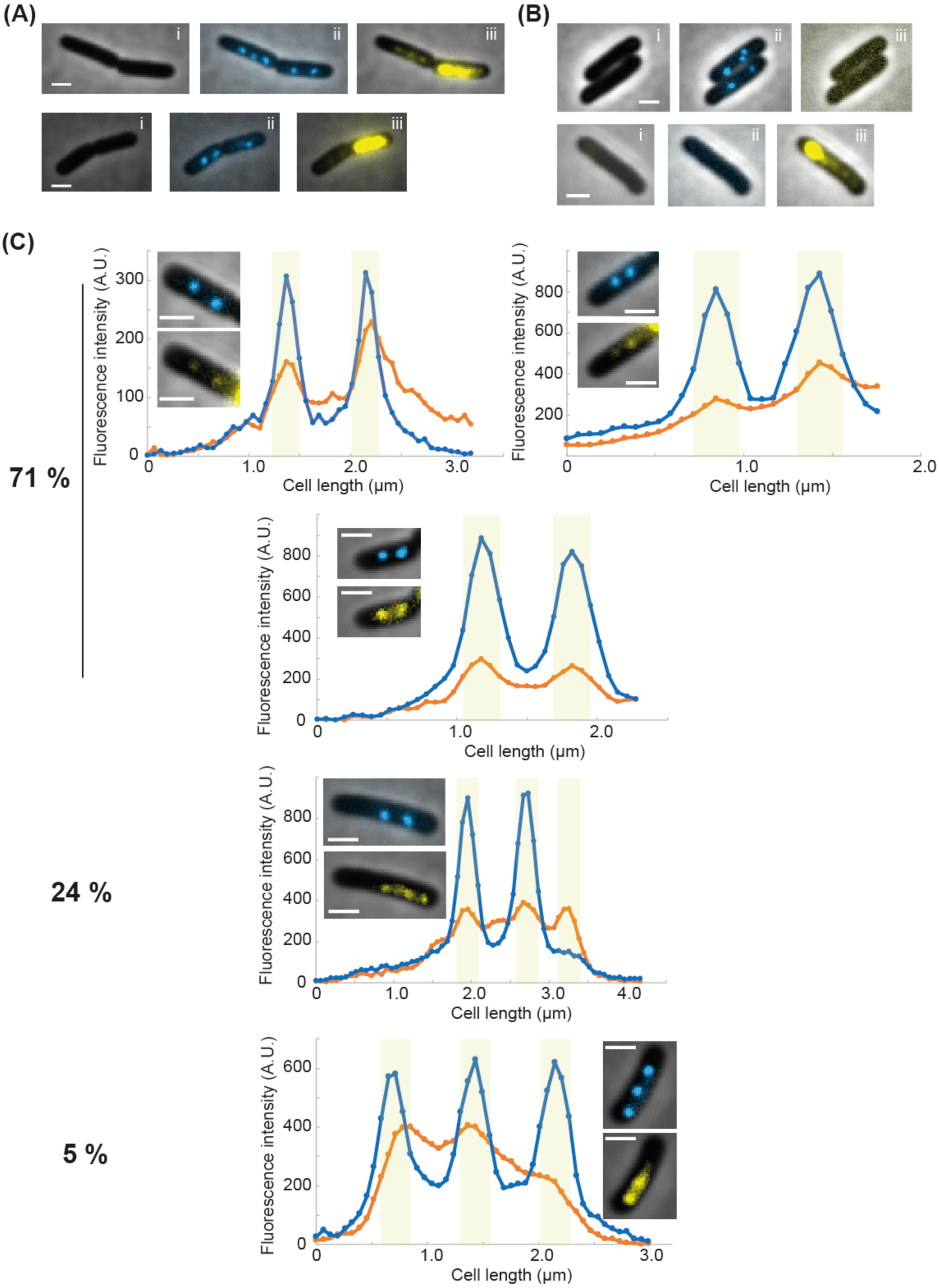
Co-localization of ParA_F_ foci and partition complexes. (A) Two typical examples of asymmetric inheritance of ParA_F_ upon cell division. Due to the oscillatory behavior of ParA_F_, most often one cell inherits most of ParA_F_ while the other a much smaller amount. Dividing cells are observed in phase contrast (i) and in fluorescence microcopy to observe ParB_F_-mTq2 (ii, blue channel) or ParA_F_-mVenus (iii, yellow channel) in overlay with phase contrast. Cells were grown at 30°C in MGlyC. (B) Fluorescence signals are specifically detected without spreading in other channels. Strains carrying mini-F plasmids expressing either only ParB_F_-mTurquoise (pJYB240; top) or only ParA_F_-mVenus (pJYB243, bottom) are grown and imaged as in (A). No leaky signal from mTurquoise2 or mVenus is observed in the yellow (iii) or blue (ii) channels, respectively. (C) Line scan analyses of fluorescence intensity along cell length. Blue and orange lines correspond to the blue (ParB_F_-mTq2) and yellow (ParA_F_-mVenus) channels, respectively. The corresponding cell images is displayed in the graph as in (A). Over 58 cells, 41 (71%) displayed the same number of ParA and ParB foci, 14 (24%) displayed more ParA than ParB foci, and 3 (5%) displayed more ParB than ParA foci. The light green area corresponds to the limit of resolution of the microscope (i.e. 4 pixels (262 nm) around the PC peaks). Scale bar: 1 μm in all images.

**Fig. 5 – Figure supplement 4.**
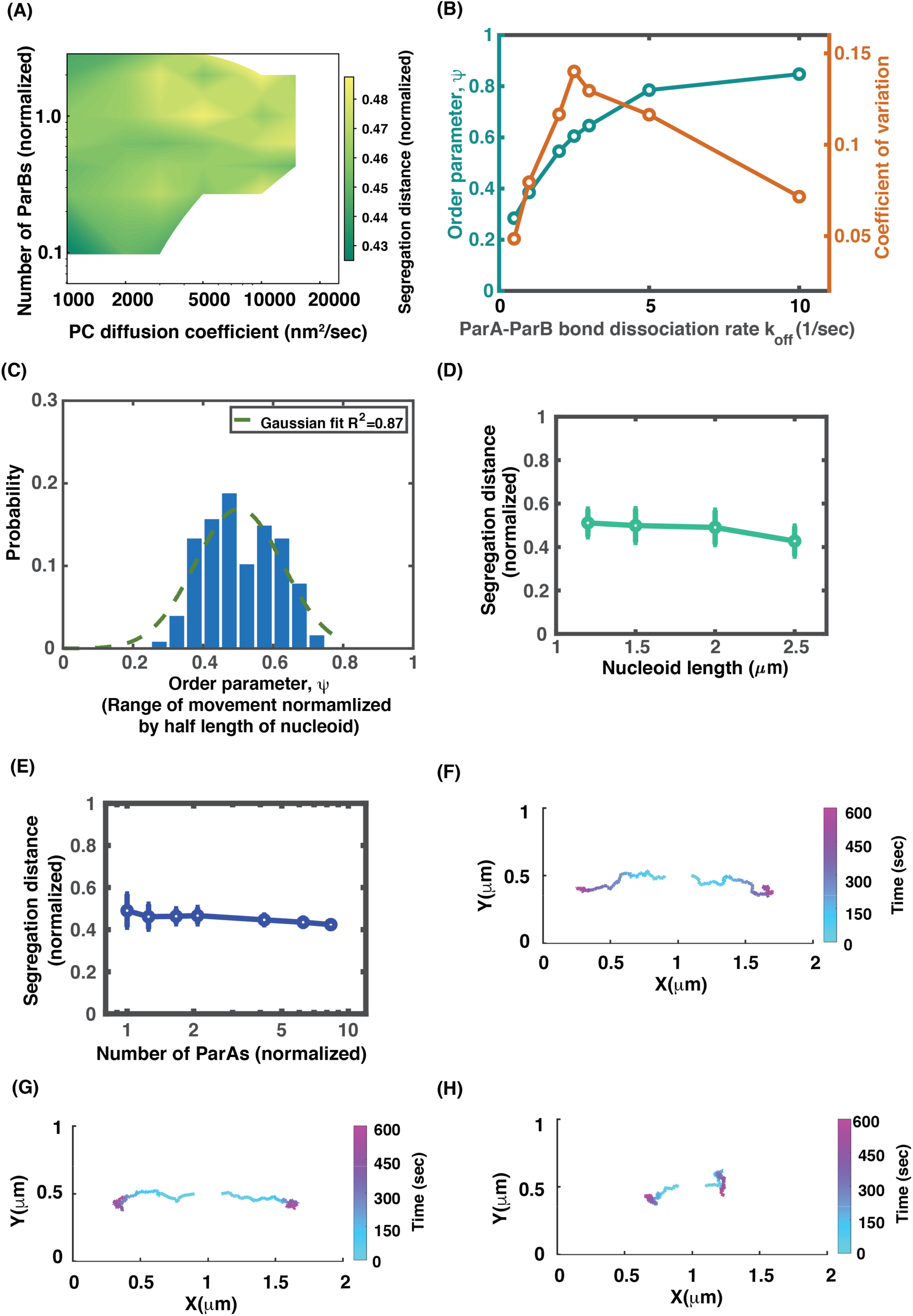
Model exploration of potential impacts of PC moving inside nucleoid on PC partition. The explored effects include the combination of a slower PC diffusion and variation in the number of PC-bound ParB available for ParA binding (A-E), and non-uniform nucleoid DNA distribution (F-H). Here, all the model calculations are performed with the full model that depicts the PC-localization of ParA. (A) Computed phase diagram of the dependence of PC segregation on PC diffusion coefficients and the number of PC-bound ParBs that is available for ParA binding. (B) Characteristics of the near-critical-point operation. (C) Statistical distribution of PC excursion distance near the critical point. (D) The segregation distance adapts to half lengths of the nucleoid near the critical point. The parameter set of (*k*_a, plasmid_, *k*_off, plasmid_) used here is the same as that in Fig. 5B. (E) Segregation distance adaptation buffers against the variations in the ParA level. For (B-E), the PC diffusion coefficient is chosen to be 2000 nm^2^/sec, 5 times slower than that in Figure 5. (F-H) Non-uniform nucleoid DNA density directs PC movement. PCs move on the nucleoid substrate surface with non-uniform DNA density along the x-direction. Three cases were simulated: (F) the density of nucleoid DNA increases from the center to the poles; (G) the density of nucleoid DNA increases from the center to the quarter positions; and (H) the density of nucleoid DNA decreases from the center to the poles. A typical simulated trajectory is shown for each case, and the normalized segregation distances averaged over 36 independent trajectories are (F) 0.69, (G) 0.63 and (H) 0.27, respectively.

